# Disentangling Stimulus & Population Dynamics in Mouse V1: Orthogonal Subspace Decomposition for Neural Representation

**DOI:** 10.1101/2025.11.10.687558

**Authors:** Nikolaos Tzanakis, Alexandros Barberis, Mario Alexios Savaglio, Ioanna Chourdaki, Stelios Manolis Smirnakis, Maria Papadopouli

## Abstract

Understanding how the primary visual cortex of mice represents the external sensory input separately from the internal states is a fundamental challenge in systems neuroscience. Our work contributes to the problem of decoupling the stimulus-driven and internally generated components of neural activity in the primary visual cortex. Internally generated (or intrinsic) activity refers to neural dynamics that are not directly driven by sensory stimuli, reflecting the brain’s ongoing, endogenous processes. Neuronal activity encodes both external stimuli and internal cortical states. The internally modulated activity, though not directly observable, can be inferred from the shared structure in population responses, and thus, serves as a proxy for the internal cortical state.

We developed a two-phase Partial Least Squares Regression (PLSR) framework that decomposes neural activity into two orthogonal low-dimensional subspaces: (1) a “population” sub-space capturing global variability shared across neurons, and (2) a “stimulus” subspace containing dimensions that discriminate between stimulus conditions while being linearly uncorrelated with the population subspace. We focus on the granular (L4) and supragranular (L2/3) layers of awake mice exposed to visual stimuli consisting of optical flow directions, using mesoscopic two-photon calcium imaging. In both L4 and L2/3 layers, many components individually yield above-chance decoding accuracy, yet a small low-dimensional subspace preserves nearly the full decoding performance of the high-dimensional population. Stimulus-driven components exhibit strong cross-mouse correlations, indicating a conserved coding scheme present in both L4 and L2/3. These components are stable across the entire recording session, reflecting robustness of the underlying representation over time. Removing the global modulation did not abolish stimulus discriminability in either layer, suggesting that information about stimulus direction is not dependent on this global signal. Both L4 and L2/3 stimulus components exhibit comparable decoding performance as well as similar tuning representations, suggesting common encoding of stimulus direction across layers.

## I. Introduction

The nervous system builds internal sensory representations through coordinated patterns of neural activation. These activity patterns simultaneously encode non-sensory variables— such as internal state, behavioral context, or developmental circuit dynamics—that modulate sensory-guided behavior [1], [2]. Ongoing endogenous processes continuously shape evoked neural responses, making the decoding of embedded representations from large-scale neural recordings a cornerstone problem in systems neuroscience. Traditionally, this challenge is tackled by mapping stimulus parameters to the magnitude of neural responses. Yet the neural activity elicited by identical stimuli is highly variable across repeated trials [3]. A contributor to this variability is internally driven activity (IDA)—intrinsic activity not directly driven by the stimulus. Once considered noise, IDA is now recognized as reflecting structured internal dynamics, including behavioral state [4]–[8] or predictive coding processes [9]. For example, rapid arousal-state transitions can modulate sensory processing [10], [11]. Stringer *et al*. [7] showed that a large fraction of trial-to-trial variability in evoked responses arises from stimulus-independent sources, including internal-state fluctuations and behavior.

A central challenge in systems neuroscience lies in extracting low-dimensional, interpretable representations from large-scale neural recordings. This requires disentangling the complex, high-dimensional neural activity to reveal the underlying structure associated with external stimuli, internal states, and behavior. Recent advances in large-scale neural recording technologies, such as two-photon calcium imaging and high-density electrophysiology (e.g., Neuropixels), now enable simultaneous measurement of thousands of neurons across multiple brain regions during diverse behavioral conditions [7], [12]–[20]. These datasets have been analyzed under both stimulus-driven and spontaneous conditions, often in awake animals exposed to naturalistic [7], [15], [17], [19] or passive visual stimuli [7], [12], [14], [16], [18], and across species including mice, monkeys, and zebrafish.

Prior work has taken several general directions to characterize the structure in such high-dimensional neural activity: (i) identifying ensembles of neurons with coordinated firing [21]–[24]; (ii) covariance-based approaches, such as PCA, SVD, and NMF [7], [19], [20], to extract dominant variance modes without directly leveraging stimulus information; (iii) jointly modeling stimulus and internal dynamics [12]; (iv) developing more expressive latent-variable models—including latent dynamical systems (LDS), variational autoencoders (VAEs), and rectified latent variable models–to build generative frameworks for modeling neural population activity, enabling the recovery of low-dimensional structure and latent factors underlying observed neuronal dynamics [13], [16]–[18], [25]; (v) studies examining how neuronal population activity, decomposed into distinct frequency components or states, relates to sensory encoding and neural variability [14], [15]. Stringer *et al*. [7] used shared-variance component analysis to identify neural dimensions accounting for activity driven by visual stimuli versus spontaneous behaviors, enabling a low-dimensional decomposition of population dynamics. They found that spontaneous behaviors—including facial movements and locomotion—drive widespread, highdimensional neural activity largely orthogonal to stimulus responses. Xia *et al*. [19] applied tensor component analysis, a multi-mode extension of PCA, in an unsupervised fashion to decompose neural activity across neurons, time, and trials, identifying trial-invariant components likely reflecting stimulus-driven responses, and trial-unique components capturing stimulus-independent variability. Carbonero *et al*. [20] used nonnegative matrix factorization (NMF) to analyze largescale calcium imaging data, showing that its positivity and linearity constraints yield interpretable components that preserve variance and capture neuronal dynamics more accurately than alternative dimensionality-reduction methods (PCA, ICA, UMAP). Triplett *et al*. [12] explicitly incorporated stimulus onset timing as a one-hot encoded regressor, convolved with an impulse-response kernel (scaled per neuron), to model evoked activity. Simultaneously, internally-generated dynamics were captured using a low-dimensional latent variable model, with both components jointly fit to disentangle stimulus-driven and spontaneous neural signals without sequential subtraction. Whiteway *et al*. [13] applied a rectified latent variable model (RLVM) to uncover hidden factors underlying cortical activity, identifying both stimulus- and behavior-related dynamics without stimulus labels. Koh *et al*. [16] developed the Calcium Imaging Linear Dynamical System (CILDS) to perform joint deconvolution and dimensionality reduction on calcium imaging data, extracting latent trajectories reflecting shared, time-varying neural dynamics while accounting for calcium decay and noise. Zhu *et al*. [17] introduced miVAE, which takes a self-supervised multi-modal approach by explicitly disentangling neural activity into stimulus-related and stimulus-unrelated latent subspaces. It jointly trains complementary generative models conditioned on both neural responses and visual stimuli, enabling the separation of stimulus-correlated structure from individual-specific variability — a clear instance of self-supervised disentanglement in population-level neural modeling. Gokcen *et al*. [18] proposed Delayed Latents Across Groups (DLAG), a dimensionality-reduction framework that disentangles bidirectional signal exchange between neural populations. DLAG separates latent activity into components relayed in each direction, identifies how these signals are represented within each population. Frequency-based approaches include Thivierge *et al*. [14], who introduced frequency-separated PCA, by applying multilinear PCA after extracting components of neural activity within distinct frequency bands to isolate temporal-scale-specific patterns. Akella *et al*. [15] developed a state-conditioned encoding framework that uses hidden Markov models of local field potentials to identify oscillatory brain states and partition neuronal variability into contributions from internal dynamics, behavior, and visual stimuli. These studies differ in the level of supervision, the sources of variance modeled, and whether stimulus and internal dynamics are fit jointly or sequentially. Some impose orthogonality constraints to enforce distinct variance sources [7], [14], [16], [19], while others allow overlapping modes. Experimental settings—including stimulus type (natural scenes vs. gratings), imaging tools, and animal state—also influence the observed dynamics and choice of methodology to be applied.

Our approach first removes components reflecting global modulation and then extracts mutually orthogonal low-dimensional subspaces from the residual activity that discriminate between stimulus directions. The same analysis is applied independently to L4 and L2/3 populations to assess laminar differences in encoding. Our work employs supervised latent-structure models designed to separate stimulus-driven from populationwide variability.

We applied Partial Least Squares Regression (PLSR) [26]–[28] within a two-phase analysis framework, a widely used approach in neuroscience for identifying low-dimensional projections of neural activity that optimally predict task-relevant variables. Rumyantsev *et al*. [29], for example, applied PLSR to two-photon recordings in mouse V1 and L2/3 pyramidal neurons to dissociate responses to distinct visual stimuli, showing that stimulus-specific dimensions are largely orthogonal to dominant noise modes. In our framework, the first phase extracts components capturing global activity patterns, likely reflecting ongoing brain states or recurrent dynamics. These projections are then subtracted from the original activity to obtain residuals. In the second phase, stimulus-discriminative components are derived from these residuals using stimulus labels. This two-stage process isolates stimulus-driven subspaces independently of global variability. Unlike purely covariance-based methods (e.g., PCA, NMF), PLSR leverages supervision to guide component extraction while preserving interpretability through linear projections. Orthogonality constraints ensure that each axis captures a distinct source of variance, enhancing interpretability and separating underlying dynamics. Finally, we introduce an iterative, cross-validated PLSR procedure that disentangles neural activity into two mutually orthogonal subspaces: one capturing global population-wide variability and another capturing stimulus-discriminative responses. Experiments consisted of continuous recordings of neuronal activity during optical flow stimulation (Monet movie) without interleaved blanks; short consecutive unlabeled segments were excluded from analysis.

**Fig. 1:**
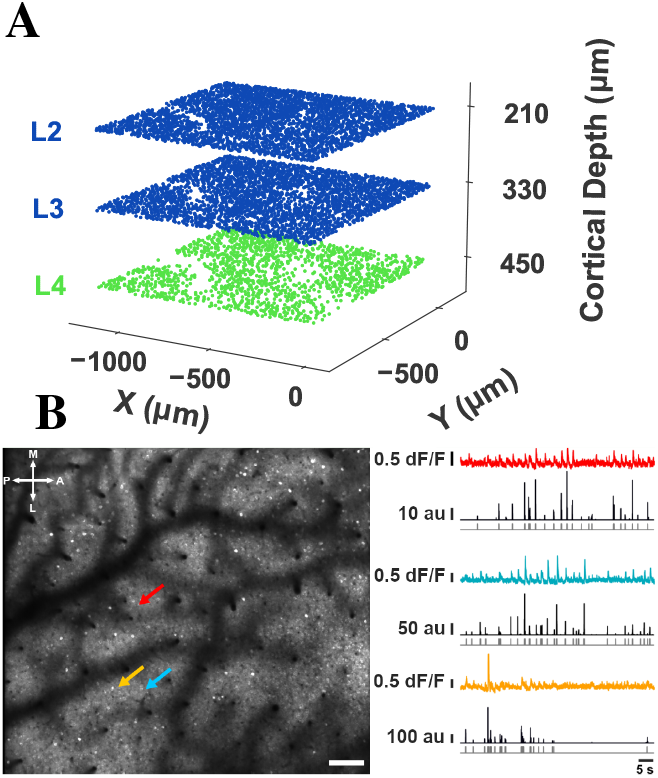
Imaging Paradigm. **A:** Illustration of L2, L3, L4 fields of view (FOVs) simultaneously acquired at 6.3Hz. L2/3: blue. L4: green. **B:** Example FOV acquired in L2/3 at depth 210 *µ*m. A: anterior, L: lateral, P: posterior, M: medial. Bar = 75 *µ*m. Color arrows indicate 3 example cell bodies whose traces are shown in color on the right. dF/F: fractional fluorescence change. au: the relative probability of firing in arbitrary units. Deconvolved firing probability traces shown in black below were obtained using the CaImAn algorithm [30], then thresholded to yield calcium “eventograms” that were analyzed. In what follows, we chose the threshold yielding population calcium event rates close to those reported in the literature [31], but results were robust to the choice of threshold. Gray traces at the bottom represent the thresholded, binarized, probability that specific imaging frames contain a calcium event (0: no event; 1: event).

Section II overviews the experiments, data collection, and preprocessing. Section III describes the PLSR-based methodology while Section IV presents performance analysis. Section V discusses our main results and outlines future directions.

## II. Experiments, Data Collection, and Pre-processing

This work focuses on data obtained from the granular (L4) and supragranular (L2/3) layers in the primary visual cortex^1^ of *five adult mice*. For each mouse, we retained approximately *60-minute neuronal recordings*, during which mice were presented with stimuli videos of smoothened Gaussian noise with coherent orientation and motion (example frame in Fig. 2), consisting of waves with **16 distinct** randomly shuffled directions of motion [32]. All 16 distinct directions of motion were presented in random order in the course of a 15-sec video. Each portion of the video corresponding to a *fixed stimulus direction* is referred to as a **segment** and lasts 937.5 ms. In the following video, the 16 directions were presented in a *different* order. 240 such videos were shown *consecutively*. The two-photon (2P) imaging recordings were preprocessed for motion correction and underwent automatic segmentation, deconvolution, and appropriate thresholding [30] (see methods in Appendix) to produce calcium “*eventograms*” that were analyzed. For each neuron, the calcium eventogram was used to compute the **calcium event rate per segment** (*ERPS*), based on 2P frames that coincided temporally with each Monet segment.

**Fig. 2:**
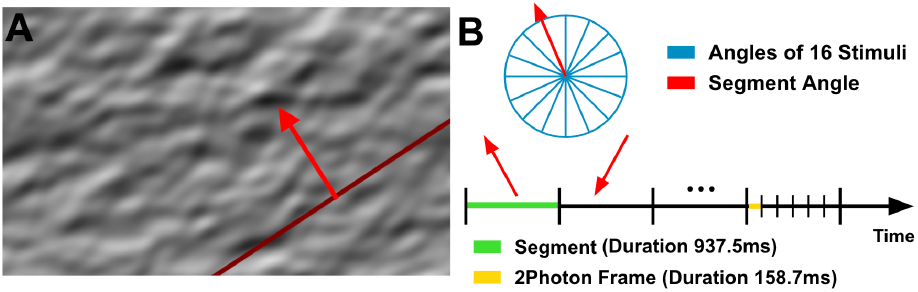
Stimulus Presentation. **A**. Example frame of “Monet” video, consisting of waves with 16 distinct randomly shuffled directions of motion, presented to the mice (i.e., stimulus) (See Appendix for additional information). The red arrow indicates the direction of motion of the stimulus **B**. Example of a sequence of segments, each segment with a fixed stimulus direction.

## III. 2-phase Partial Least Squares Methodology

We employ a two-phase Partial Least Squares Regression (PLSR)-based approach that aims to explicitly separate stimulus-driven from population-related neural variability. This procedure yields two low-dimensional, mutually orthogonal subspaces: **1)** a population subspace that captures population dynamics, and **2)** a stimulus subspace that yields a representation in which neural responses to different stimulus classes can be distinguished and do not linearly account for the population subspace that is identified in 1);

### Phase 1: Identification of Population Activity Components

Let **X** ∈ ℝ^*N×T*^ denote the event rate per segment, where *T* is the number of time segments and *N* is the number of neurons. PLS Regression constructs a sequence of components, *iteratively*, identifying directions in the input data that maximize covariance with the target variable. Specifically, the first component is the linear projection of **X** (via weights *w*_1_) that maximizes its covariance with a corresponding projection of aggregate population activity (*Y*_*p*_) which is defined at each time t as the sum of single-neuron event rates, *Y*_*p*_(*t*) =∑ _*i*∈𝒩_ *r*_*i*_(*t*), where *r*_*i*_(*t*) is the event rate of neuron i at time t, and 𝒩 is the set of neurons. In this work, we use the term global modulation (or population activity) to refer to fluctuations in neural activity that are shared across a large fraction of neurons within a cortical layer. This aggregate captures widespread co-activation patterns that likely reflect internal state changes, arousal fluctuations, or recurrent network dynamics. When analysis is restricted to a specific layer (L4 or L2/3), 𝒩 includes *only* the neurons of that layer. Once this component is extracted via deflation, the residuals in both **X** and **Y**_**p**_ are updated, and the next component is computed *iteratively* using the updated residuals. This procedure continues until the identification of *D*_1_ components, yielding the following: 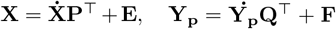, where: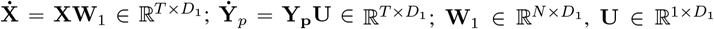; and residuals **E, F**.

**TABLE I:**
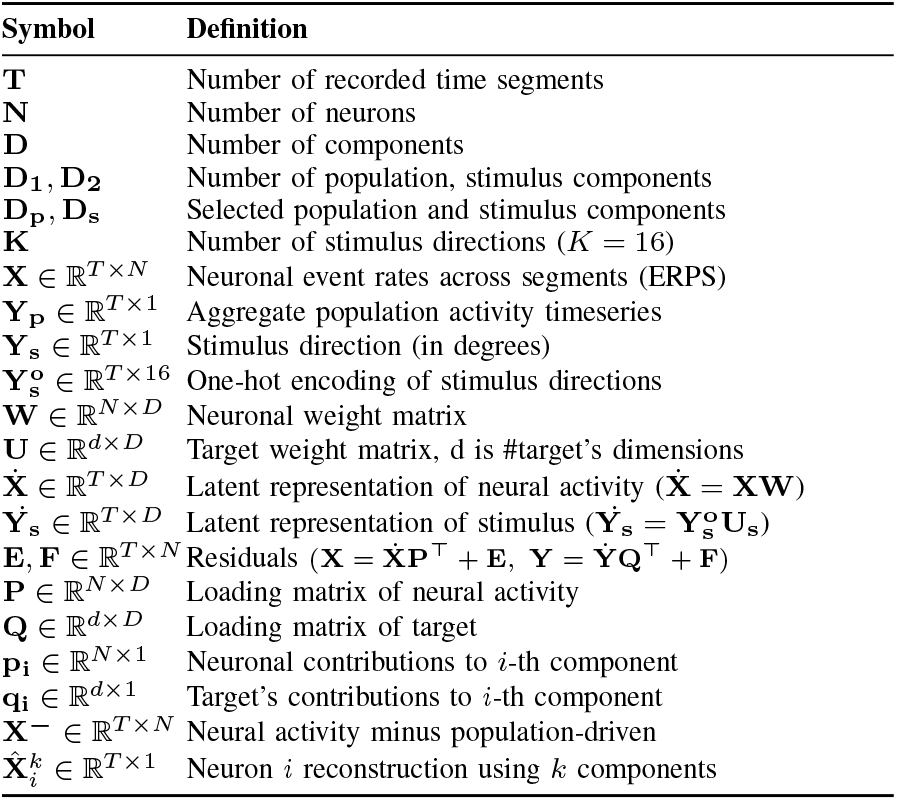
Definition of Variables.

#### ALGORITHM I

***Partial Least Squares*** *Regression (PLS) Phase 1: (***X**_**0**_ = **X, Y** = **Y**_**p**_*); Phase 2:* 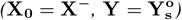

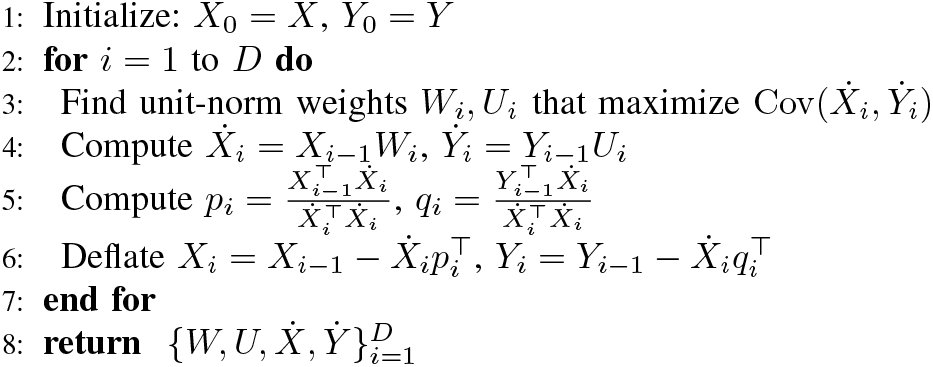

### Phase 2: Identification of Stimulus Components

After selecting the population-driven components *D*_*p*_ consisting of the first PLS components of Phase 1 that explain the aggregate population activity (**Y**_**p**_), the residual activity (**X**^**−**^) is calculated as follows: 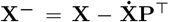where 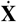 and **P** correspond to the population-driven components *D*_*p*_ and their associated loadings, respectively. **X**^**−**^ serves as the input to a second PLS regression run, with target the stimulus 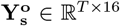, which is defined as: Let 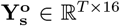 be the stimulus direction variable **Y**_**s**_ in one hot encoding. One-hot encoding is a method for representing categorical variables as binary vectors. In this setting, the stimulus direction variable **Y**_**s**_, which takes one of 16 discrete values at each time segment, is encoded using one-hot encoding to produce a matrix 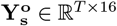. Each row of 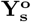 contains a *single* 1 indicating the stimulus direction at the corresponding time segment, with all other entries being 0. PLS regression is then applied to extract mutually orthogonal latent dimensions of the residual neural activity **X**^**−**^ that maximally covary with the stimulus direction matrix 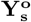, thereby isolating components of residual activity most informative for discriminating stimulus directions. The PLS regression extracts mutually orthogonal latent dimensions of **X**^**−**^ that covary with latent dimensions of 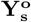, 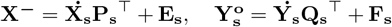. We selected the first stimulus components returned by PLS (dimensions: *D*_*s*_) whose cumulative decoding accuracy for stimulus direction reached a plateau, comparable to the performance obtained when using the entire neural population. Our analysis shows that, in general, 6 to 8 stimulus components are sufficient to achieve decoding accuracy comparable to that obtained with the full neural population (Fig. 3I). First Stimulus-component weights are near zero and symmetric (Appendix, Fig. 6A). The first population component (Population ID 1) exhibits a positive offset (Appendix, Fig. 6B), as expected, given that the target variable corresponds to the aggregate-population activity. With matched neuron counts, the weight distributions in L4 and L2/3 have similar shapes—apparent differences in Fig. 6 arise from a unit-norm PLS scaling artifact (||*W*_*i*_ ||= 1). In a heuristic manner, we take into consideration the non-linear dependencies that may arise between the population activity components and aggregate population activity, by computing the z-score-normalized mutual information (znMI, see Appendix) between each component 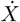 and aggregate population activity **Y**_*p*_, tested for statistical significance against a null distribution generated via permutation. Specifically, we order the population activity components based on their znMI with the aggregate population activity (Fig. 3A). We use the same heuristic to sort the stimulus components that are identified in Phase 2, in descending order with computing the z-normalized mutual information (znMI) of each stimulus component 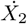 and Stimulus *Y*_*s*_ (Fig 3F). Our analysis was performed on a training set comprising 80% of randomly shuffled data segments. Model evaluation was subsequently carried out on the remaining 20% of the data, referred to as the test set, to assess generalization performance. Each neuron’s event rate was standardized by subtracting the mean and dividing by the standard deviation computed from the training set only, and these training-set statistics are applied to both the training and test sets to avoid using any information from the test set during training. We also repeated the two-phase PLS analysis in reverse order: First, we identified the stimulus components and selected those whose decoding accuracy for stimulus prediction matched that of the full population. Then, using the residual neural activity obtained after removing the selected stimulus components, we identified the population components with a second PLS analysis. However, the first stimulus components retained significant information about aggregate population activity. This asymmetry implies that large-scale population activity is not orthogonal to the sensory representation but covaries with it. Fluctuations related to internal states (e.g., arousal, movement, attention) influence neurons tuned to visual features in a correlated way.

**Fig. 3:**
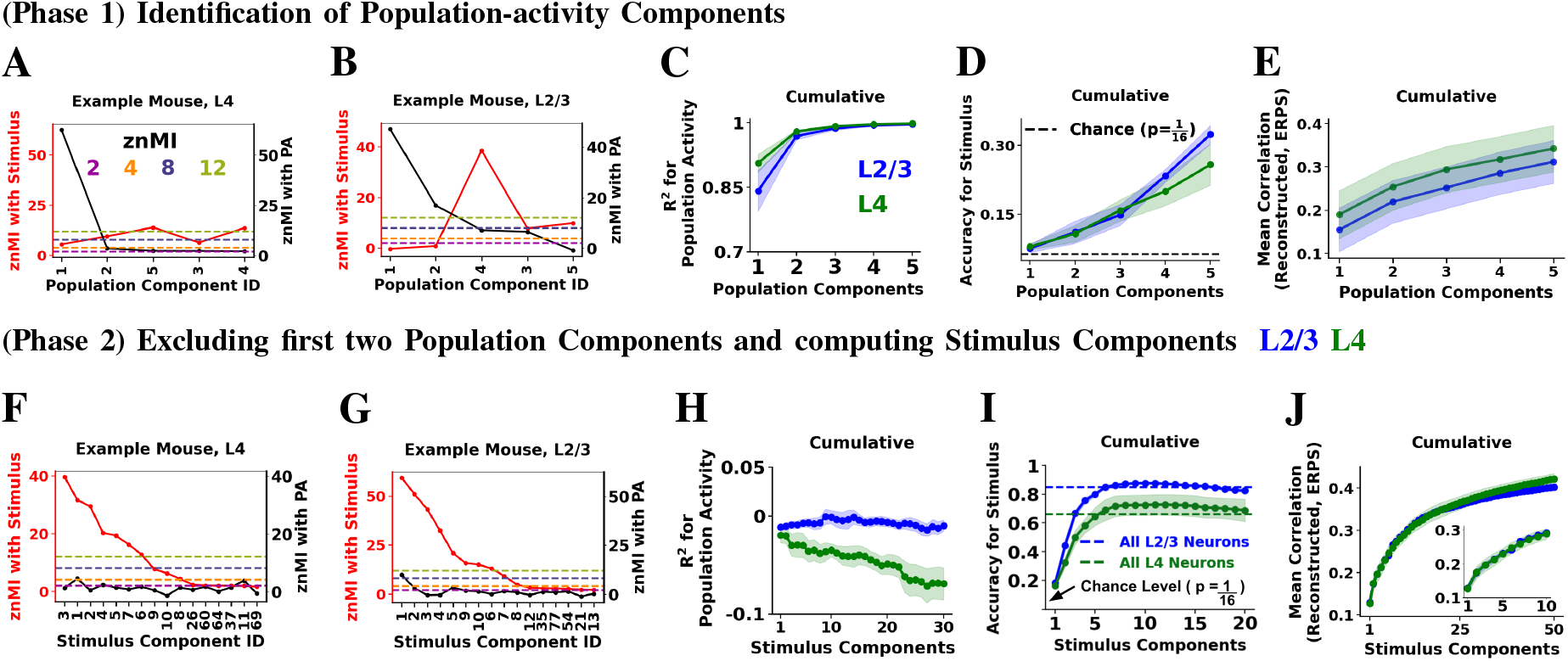
Identification of Population- and Stimulus-Driven Components. **A:** Z-score-normalized mutual information (znMI) between population components of L4 and *aggregate population activity* of L4 (black) and *stimulus direction* (red), for an example mouse. Dashed horizontal lines indicate znMI thresholds (2, 4, 8, and 12), shown in dark magenta, dark orange, dark slate blue, and dark olive-green, respectively. **B:** Same as 3A, but analysis is performed on L2/3. **C:** Coefficient of determination (R^2^) for predicting aggregate population activity as a function of the number of population components used in Linear Support Vector Regression. **D:** Decoding accuracy as a function of number of population components used for predicting the stimulus direction using Logistic Regression. **E:** Mean Pearson correlation between each neuron’s original event-rate per segment time series and its reconstructed signal as a function of population components used, averaged across all neurons in corresponding area, plotted as a function of the number of population components used (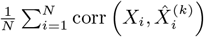), where k denotes the number of components. **F:** Z-score-normalized mutual information (znMI) between stimulus components (x-axis) and stimulus direction in red, aggregate population activity in black, analysis performed on L4 neurons. Dashed horizontal lines corresponds to different znMI thresholds, as in 3A. **G:** Same as 3F, but analysis is performed on L2/3. **H:** Coefficient of determination (R^2^) for predicting aggregate population activity as a function of the number of stimulus components used, using Linear Support Vector Regression. **I:** Cumulative stimulus decoding accuracy as a function of the number of stimulus components. Dashed horizontal lines represent the accuracy obtained when using the event rate per segment of all neurons as features (blue for L2/3 and green for L4). Black arrow indicates the chance-level accuracy (1/16) for predicting stimulus direction. **J:** Mean Pearson correlation between each neuron’s original event-rate time series (per segment) and its reconstructed signal, obtained as a function of stimulus components used. Correlations were averaged across all neurons and plotted as a function of the number of stimulus components used. Inset: cumulative correlation of the first 15 stimulus components. Lines show averages across mice (n = 5), with shaded areas indicating the standard error of the mean across mice. PLS was performed considering *D*_*p*_ = 2 population components and *D*_2_ = 100 stimulus components were generated.

In two-phase PLSR, the “orthogonality” refers to the statistical decorrelation achieved through the sequential deflation process in PLS regression, which ensures that each extracted component captures a distinct source of shared variance between neural activity and the corresponding target variable. Orthogonality here does *not* imply strict geometric independence between the stimulus and population subspaces but rather that their projections are linearly uncorrelated by construction, minimizing shared variance and enhancing interpretability. All classifiers, regressors, and the PLSRegression model are implemented using scikit-learn [33].

## IV. Analysis of the Components

The first population component (ID 1) shows a high znMI with aggregate population activity and, depending on layer and mouse, may also carry stimulus-related information (Figs. 3A, 3B). Including two components explains about 97% of the variance in aggregate population activity on the test set across all mice and layers (Fig. 3C). The population components collectively capture information about stimulus direction, with decoding accuracy remaining above chance and improving as additional components are included in both L4 and L2/3 (Fig. 3D). Because the first two population components already explain most of the aggregate population activity, subsequent PLS components are fit to residual targets that no longer reflect overall population-activity variance but instead capture more subtle, residual variability that happens to carry information about the stimulus. For each neuron, the population-driven signal is reconstructed from the first two population components and compared to its original ERPS. The population-driven signal of individual neurons correlates with their event rate per segment (ERPS), with mean correlation coefficients across neurons of *r* = 0.20 in L2/3 and *r* = 0.25 in L4 (Fig. 3E). Stimulus components computed after removing the first two population components via deflation exhibit high znMI with the stimulus, and several also retain substantial non-linear information about aggregate population activity. In every layer and mouse, there is consistently at least one stimulus component with high znMI to the stimulus and a znMI greater than 4 with the aggregate population activity (Figs. 3F, 3G). To verify that stimulus components do not exhibit linear cumulative information about aggregate population activity, we computed the *R*^2^ between stimulus components and the aggregate population activity, consistently obtaining *R*^2^≈ 0 (Fig. 3H). The low-dimensional representation of neural activity (6 dimensions for L4, 7 for L2/3) retains nearly the same predictive accuracy for stimuli direction as the full high-dimensional population. Consistent with Stringer *et al*. [34] using few components, it is sufficient to achieve classification performance comparable to that obtained using the entire population (Fig. 3I). Up to a certain point, adding more components improves accuracy; beyond that, however, their contribution becomes detrimental due to model overfitting. We reconstructed the stimulus-driven signal of each neuron and evaluate its correlation with the original event rate per segment (ERPS). The pearson correlation increases sharply with the first few components and then rises more gradually. Mean correlations are comparable between L4 and L2/3 (Fig. 3J).

Stimulus-driven components that are identified by our twophase PLSR, effectively separate neuronal activity to best capture differences in stimulus directions. The *first* onedimensional component, that exhibits the *highest covariance* with the visual stimulus direction, achieves significantly higher accuracy and information for stimulus decoding compared to the aggregate population activity (Population Activity) and a control of event rate per segment (NULL ERPS) (Fig. 4A). The first one-dimensional component shows direction tuning at 0^°^ and 180^°^. Although the precise angular position of the peaks varies somewhat across mice in both L2/3 and L4, at least three mice exhibit consistent peak locations. (Fig. 4B). The tuning for stimulus angle of the first eight stimulus components reveal a distinct structure: the leading PLS components (e.g., ID 1, ID 2), which best separate neural responses by stimulus direction, captured by the order of PLS algorithm, predominantly reflect orientation tuning, while subsequent components (e.g., IDs 3-5) capture direction selectivity. The remaining stimulus components (e.g., IDs 6–8) exhibit weaker modulation and higher frequency across stimulus angles (Fig. 4C). To evaluate whether the higher decoding accuracy observed in L2/3 compared to L4 (Fig. 3I) results from its larger neuronal population, we repeated the analysis by randomly selecting subsets of L2/3 neurons matched in number to the L4 population (30 random samples). This analysis revealed that L2/3 and L4 exhibit comparable decoding accuracy when 8 stimulus components are used (Fig. 4D). We then excluded the first eight components—which all together yield higher accuracy than the full population—and recalculated stimulus prediction accuracy using the remaining components. The residual components exhibit an accuracy of approximately 0.2 for L4, and 0.3 for L2/3 (Fig. 4E). Examining the separability of stimulus classes for each stimulus component individually, we found that the first ~ 15 components achieve above-chance accuracy (Fig. 4F). For the residual activity—neural activity after removing the first two population components—the top stimulus-driven components, which best discriminate the stimuli, also explained more variance than the remaining components, indicating that these axes capture the most prominent dimensions of the residual neural activity. Considering the explained variance on the training set, L4 consistently shows a slight increase compared to L2/3, due to the smaller size of the neurons included in L4 (Figs. 4G, 4H). When the analysis is performed using a randomly selected subset of L2/3 neurons matching the size of the L4 population, the explained variances are comparable. Across mice, the first two population components explained 6.1 ± 1.5 % of the variance in L2/3 activity and 8.0 ± 2.1 % in L4. We examined the stability of stimulus components across mice and over time (Fig. 5). After computing the components, we aligned the movie segments so that time *t* corresponded to the same stimulus angle, and then calculated pairwise absolute pearson correlations of the components throughout the recording period. The first five components show significant correlations across mice, indicating a comparable population-level stimulus code in both L4 and L2/3 (Figs. 5A, 5B). To assess temporal consistency, we divided the 60-minute experiment into consecutive 15-minute intervals and applied the two-phase PLS method to each segment separately to estimate the stimulus components within each time window. We then calculated pairwise absolute pearson correlations between components with the same IDs across periods, following the same stimulus-angle alignment procedure described above. The leading stimulus-driven components remain highly consistent across time, consistent with recent work [35] (Fig. 5C: L2/3, similar results on L4).

**Fig. 4:**
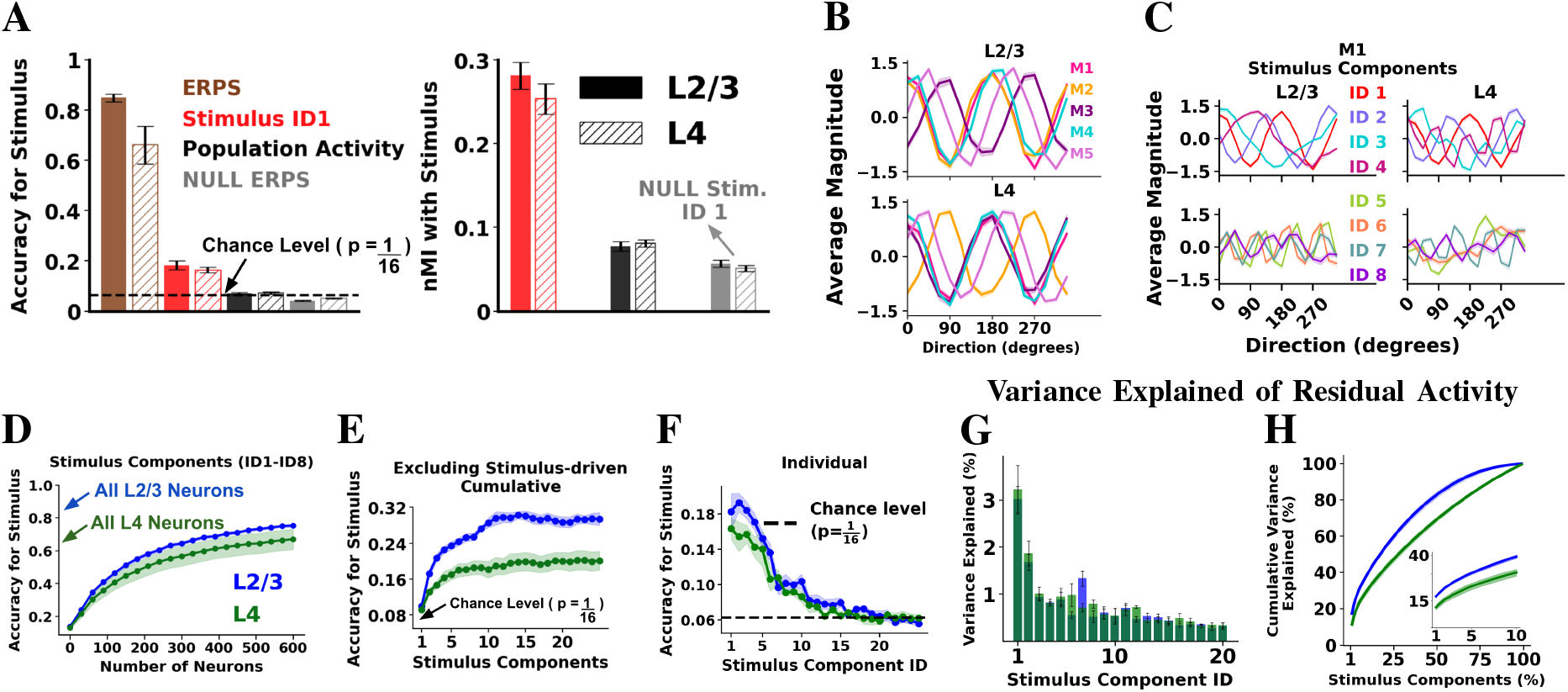
Analysis of Stimulus Components. **A:** Left Plot: Accuracy of predicting stimulus direction (*Y*_*s*_) using different inputs: Event rate per segment set (ERPS) with all neurons in the corresponding layer, first stimulus component (Stimulus ID 1), aggregate population activity of corresponding layer (Population Activity), and a circularly shift of ERPS (NULL ERPS). Right Plot: Normalized mutual information (nMI) between stimulus direction (*Y*_*s*_) and (1) Stimulus ID 1 (red), (2) Population Activity (black), and (3) null Stim ID 1 (gray). Null Stim ID 1 was computed using Two-phase PLS, with the event rate per segment of each neuron randomly circularly shifted, while keeping the observed aggregate population activity (*Y*_*s*_) as a target on Phase 1. **B:** Tuning curves of the first stimulus component (ID 1) across stimulus directions for different mice in L2/3 and L4. Components were standardized to account for differences in neuron counts across mice and layers. **C:** Tuning functions of the first eight stimulus components across stimulus directions in the test set for an example mouse, with L2/3 shown on the left and L4 on the right. **D:** Prediction accuracy for stimulus direction (y-axis) using the first eight stimulus components (ID1–ID8) derived from varying numbers of neurons (x-axis). Neurons were randomly sampled 30 times for each population size. “All L2/3 neurons” indicates the prediction accuracy of stimulus direction using as features the ERPS of all L2/3 neurons. The same analysis was performed for L4 neurons. **E:** Prediction accuracy for stimulus direction as a function of the remaining stimulus components, after excluding the first eight stimulus-driven components (ID 1–ID 8). The PLS analysis was performed on the entire population of L2/3 and L4 neurons. **F:** Accuracy of predicting stimulus direction using a single stimulus component, across mice, evaluated with Logistic Regression. **G:** Variance explained of residual activity (**X**^−^ by each stimulus component in the training set. **H:** Cumulative variance explained for the residual activity as a function of the percentage of stimuli components used. Inset plot focuses on the first 10% of stimuli components. Shaded regions and error bars indicate the standard error of the mean across mice (n=5).

**Fig. 5:**
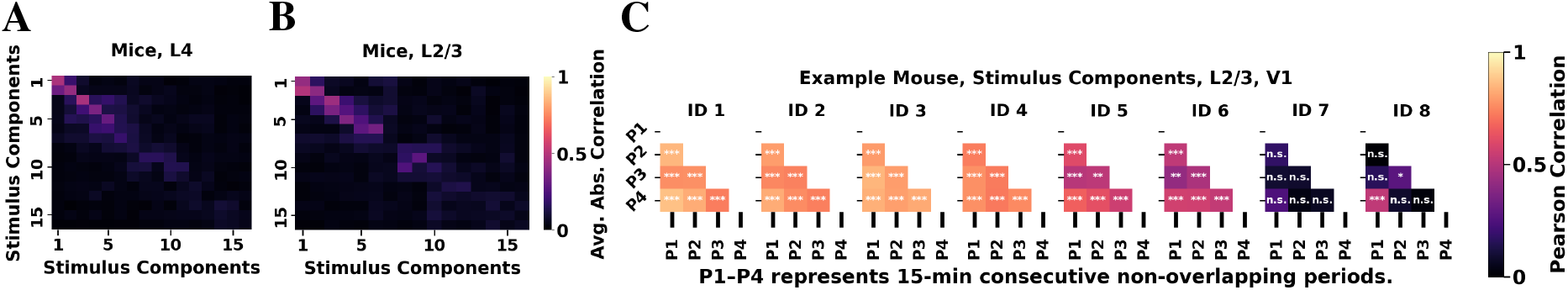
Stimulus-driven Representations Show Stable Coding Across Mice and Time. **A:** Absolute Pearson correlation between stimulus Component i of mouse *M*_*a*_ and stimulus Component j of mouse *M*_*b*_ computed for all possible pairs of mice (5 mice; 10 pairs) in entire L4. Correlations were averaged across all 10 mouse pairs for each pair of stimulus components. **B:** Same as in A, but for entire L2/3. For a fair comparison with L4, we randomly selected same number of L2/3 neurons as in L4. This sub-sampling was repeated 30 times, and the resulting correlation matrices were averaged across runs. Because the presentation order of stimulus angles differs across mice, we first computed the stimulus components for each mouse using its own aligned stimulus-angle order. We then aligned each T x 1 Stimulus component time series by reordering each 16 x 1 consecutive movie segment, so that the stimulus presentation sequence began at 0, followed by 22.5, and so on up to 337.5. **C:** Absolute Pearson Correlation across different recording periods for the same stimulus component, identified as ID k, for the first 8 Stimulus components in L2/3, for an example mouse. The two-phase PLS analysis was performed independently for each 15-minute consecutive recording period, and correlations were then computed between the corresponding components (same ID) across periods. Statistical significance was assessed using a parametric one-sided *z*-test: for each correlation cell, the corresponding values from 100 control correlation matrices were used to compute a null mean (*µ*) and standard deviation (*σ*). The observed correlation (*r*_obs_) was compared to *µ* using 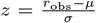 and a one-tailed *p*-value was obtained from the standard normal distribution. A small *p*-value indicates that the observed correlation is statistically greater from the control distribution. Stars mark significance thresholds (^∗^*p <* 0.05; ^∗∗^*p <* 0.01; ^∗∗∗^*p <* 0.001; n.s., not significant).

**Fig. 6:**
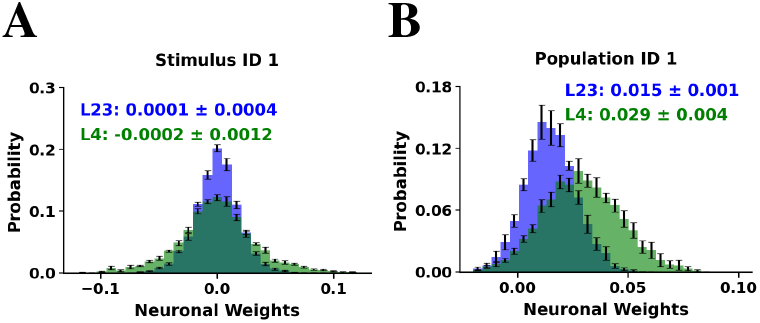
Weights of the first component for each neuron. **A:** Acrossmice distributions of neuronal weights (*W*) for Stimulus component ID1: L4 (green) and L2/3 (blue). **B:** Same as A but for the Population component with ID 1. The mean ± standard deviation of the sample means across mice (n=5) are reported in the histograms’ insets, while error bars correspond to the standard error of the mean (SEM) across mice.

## V. Conclusions and Future work

We introduced a two-phase, orthogonal PLSR decomposition that disentangles global population dynamics from stimulus-driven activity in mouse V1. This framework isolates a low-dimensional stimulus subspace that retains nearly the full decoding power of the high-dimensional population, remains stable over time, and is conserved across animals. The stimuli representations in L2/3 and L4 are largely comparable, with both layers exhibiting similar tuning properties and decoding power for the stimulus. Importantly, stimulus discriminability persists after removing global modulation, demonstrating that the V1 stimulus code is not carried by the global signal but instead by an orthogonal, compact representation. Although limited by its linearity and lack of temporal dynamics, these constraints enhance the interpretability of the decomposition. Looking forward, combining this frame-work with nonlinear and temporally structured models—such as those that partition time-dependent latent variables into “content” (e.g., angle, luminance) and “style” (e.g., internal state idiosyncrasies) using multi-modal identifiable variational autoencoders [17], [36]—offers a promising path toward richer characterizations of cortical coding. By showing that stable, low-dimensional, and layer-specific stimulus representations can be recovered even after controlling for global fluctuations, our work provides both a methodological tool and a conceptual advance for disentangling external and internal influences in large-scale neural activity.

## Acknowledgments

We would like to thank the members of Tolias Lab at the Department of Neuroscience at Baylor College of Medicine in Houston, Texas for performing the two-photon (mesoscope) experiments, sharing the datasets with us, and providing valuable feedback about the data. We are also grateful to Ganna Palagina for preprocessing the data and developing the data pipeline, and to Ioannis Smyrnakis and Georgios Keliris for insightful discussions on various aspects of the analysis. This work has received funding from the European Union’s Horizon 2020 research and innovation program under the Marie Skłodowska-Curie grant agreement No 101007926 as well as from the Hellenic Foundation Research Institute (HFRI) with the neuron-AD project number 04058 and neuronXnet project number 2285 (PI: Maria Papadopouli). It has been partially supported by project MIS 5154714 of the National Recovery and Resilience Plan Greece 2.0 funded by the European Union under the NextGenerationEU Program. Finally, this research was also supported by R01 NS113890, and R21 NS127299 (PI: Stelios Smirnakis). AWS resources were provided by the National Infrastructures for Research and Technology GRNET and funded by the EU Recovery and Resiliency Facility.

## Appendix

### Mouse Lines and Surgery

Five adult mice (10-12 weeks of age), expressing GCaMP6s in excitatory neurons via SLC17a7-Cre and Ai162 transgenic lines, were anesthetized and a 5mm craniotomy was placed over visual cortex as described [37]. Each mouse recovered for ~ 2 weeks prior to the first experimental imaging session.

### Experimental Data Collection

The animals underwent mesoscopic two-photon imaging covering most of dorsal area V1 and nearby extrastriate cortex, while being head-fixed on a treadmill in quiet wakefulness. Images were acquired at 6.30072 Hz over a ~1.2×1.2 mm^2^ field of view sampling simultaneously across 4 planes corresponding to V1 layers 2 (80-210 mm), 3 (285-330 mm), 4 (400-450 mm) and 5 (500 mm). Images were preprocessed in standard fashion for motion correction and underwent automatic segmentation and deconvolution using the CNMF CaImAn algorithm [30]. The deconvolved signal was thresholded appropriately to yield calcium “eventograms” that were used for analysis. The threshold yielding calcium event rates closer to those reported in the literature [31] was selected. Neurons located less than 15mm from the periphery of the field of view (FOV) were excluded in order to avoid potential edge effects arising from incomplete correction of motion artifacts.

### Monitor Positioning and Retinotopy

Visual stimuli were presented to the left eye with a 31.1×55.3cm^2^ (h×w) monitor (resolution of 1440×2560 pixels) positioned 15cm away from the mouse eye. Pixelwise responses across a 2400×2400 *µ*m^2^ to 3000×3000 *µ*m^2^ region of interest (0.2 px/*µ*m) at 200-220*µ*m depth from the cortical surface to drifting bar stimuli were used to generate a sign map for delineating visual areas [38]. The directional trial response was measured by taking the difference in cumulative deconvolved activity at the linearly interpolated trial onset and offset time points. Trial responses per direction were modeled as a two-peak scaled von Mises function ([37]). The two peaks share a preferred orientation, baseline, and width, but their amplitudes are fit independently. This function was fitted to minimize the mean squared error of all trial responses *across 16 directions* using the L-BFGS-B optimization algorithm [39]. Significance and goodness of fit were calculated by permutation. Trial direction labels were randomly shuffled among all trials for 1000 refits. The goodness of fit was calculated as the difference in *fraction variable explained (FVE)* between the original fit FVE and the median FVE across all 1000 shuffled fits. The p-value was calculated as the fraction of shuffled fits with a higher FVE than the original fit.

### Directional Visual Stimulus

A stimulus using smoothened Gaussian noise with coherent orientation and motion was presented to the mice. An independently identically distributed (i.i.d.) Gaussian noise movie was passed through a temporal low-pass Hamming filter (4Hz) and a 2-d Gaussian filter (*σ* = 4.4 at the nearest point on the monitor to the mouse). Each scan contained 72 blocks, with each 15-second block comprising 16 equally distributed and randomly ordered unique directions of motion between 0-360 degrees with a velocity of 42 degrees/s at the nearest point on the monitor. An orientation bias perpendicular to the direction of movement was imposed by applying a bandpass Hanning filter *G*(; *c*) where is the difference between the image 2d Fourier transform polar coordinates *ϕ* and trial direction *θ*, and

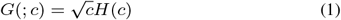

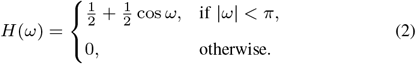

Here, c = 2.5 is an orientation selectivity coefficient. The resulting kernel is 72 full width at half maximum.

### Z-scored normalized Mutual Information (znMI)

The mutual information (MI) between two random variables quantifies the amount of information one variable contains about the other, i.e., it measures the *reduction in uncertainty* about one variable *when the value of another one is known*. Therefore, the mutual information *MI*(*X*; *Y*) between two jointly discrete random variables *X* and *Y*, with individual states *x* ∈ *X* and *y* ∈ *Y*, respectively, can be defined as: 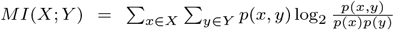 where *p*(*x, y*) is the joint probability density distribution function of *X* and *Y*, and *p*(*x*) and *p*(*y*) are the marginal probability distributions of *X* and *Y*, respectively. Thus, the *normalized mutual information (nMI)* between two jointly discrete random variables *X* and *Y* is defined as: 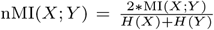 where *MI*(*X*; *Y*) is the mutual information between variables (*X*) and (*Y*), *H*(*X*) is the entropy of *X*, and *H*(*Y*) is the entropy of *Y*. When Y represents the stimulus *Y*_*s*_, we normalize the mutual information solely by the stimulus entropy **Y**_**s**_. This produces a more intuitive interpretation, as it can be understood as the proportion of the stimulus entropy explained by the neuron activity. To determine the significance of the nMI between a variable X and a variable Y, we compare it with that obtained from a control (null) distribution. More specifically, for each variable *i*, we randomly circularly shift its original time series *X*_*i*_ and then estimate the nMI with the time series *Y* (i.e.,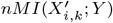)), for the k-th circularly shifted instance 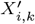 of *X*_*i*_. The above is repeated *K* = 100 times to obtain the control values of variable *i* (for *k* = 1, …, *K*). We then compute the *z-score* mutual information *zMI*(*X*_*i*_) of variable *i* as: 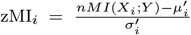 where *nMI*_*i*_ is the normalized mutual information between the observed (actual) *X*_*i*_ of variable *i* with *Y*, 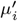 is the average normalized mutual information of the *K* control values (i.e., 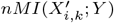)), and *σ*_*i*_ is their standard deviation. To discretize a continuous distribution (e.g., aggregate population activity or a component) so we can calculate mutual information, we use Knuth’s Bayesian optimal binning method [40], which determines the number and width of bins by maximizing a Bayesian fitness function, resulting in a piecewise constant approximation of the joint and marginal probability distributions.

## Footnotes

1 V1 receives sensory inputs in layer 4 (L4) processed vertically through the cortical column and laterally within each layer and then projected “forward” to higher areas by V1 layers 2/3 (L2/3), and “backwards” as feedback to lower areas by V1 L5/6.

